# A targeted drug-repurposing strategy identifies Tavaborole *(Kerydin)* as a potent fungistatic agent against *Candida auris*

**DOI:** 10.64898/2026.02.27.708474

**Authors:** Rounik Mazumdar, Ana Bjelanovic

**Affiliations:** Max Perutz Labs Vienna, Vienna BioCenter, Center for Medical Biochemistry, Medical University of Vienna, Vienna, Austria

**Keywords:** *<u>Keywords</u>*: *Candida auris*, Antifungal resistance, AMR, Drug-repurposing, Tavaborole, Amphotericin B, Proteomics, Electron microscopy

## Abstract

*Candidozyma auris* (*Candida auris)* is an emerging multidrug-resistant fungal pathogen posing a major global health threat. In this study, we employed a targeted drug-repurposing strategy to identify novel indications for existing FDA-approved compounds against *C. auris*, leading to the identification of Tavaborole as a potent fungistatic agent. Tavaborole displayed robust activity across all five tested clades of *C. auris*, as well as against *Candida albicans* and *Candida glabrata*. To investigate drug resistance mechanisms of *C. auris*, we applied quantitative proteomics analyses following exposure to Tavaborole and Amphotericin B (AmB), complemented by electron microscopy. Proteomic profiling revealed that *C. auris* mounts distinct but overlapping adaptive responses to antifungal stress, involving stress response pathways, metabolic reprogramming and amino acid biosynthesis. While Tavaborole primarily induced targeted stress adaptation, AmB triggered a broader, multi-pronged resistance response including oxidative stress mitigation, osmolyte production and metabolic remodeling. Shared alterations in glycogen metabolism and amino acid biosynthesis suggest conserved antifungal adaptation mechanisms. Altogether, this study highlights Tavaborole as a promising antifungal candidate against *C. auris,* sheds novel insights into drug resistance mechanisms employed the pathogen and delivers a drug-repurposing procedure highly customizable to target other microorganisms.

**Importance:** *Candida auris* is an emerging multidrug-resistant fungal pathogen responsible for healthcare-associated infections representing a high-priority antimicrobial resistance (AMR) threat due to its limited treatment options, high transmissibility, and capacity to cause severe and often fatal outbreaks. The slow pace of antifungal drug development underscores the urgent need for alternative strategies to expand the antifungal arsenal against priority pathogens such as *C. auris*. In this study, we demonstrate that a targeted drug-repurposing approach can efficiently identify antifungal activity from a small, curated set of FDA-approved compounds, leading to the discovery of Tavaborole as a fungistatic agent with broad activity across multiple *C. auris* clades. By integrating a customizable drug screening procedure with quantitative proteomics and electron microscopy, this work provides insights into antifungal resistance mechanisms. This study highlights how rational drug-repurposing strategies can rapidly identify clinically relevant drug candidates to counter emerging pathogens and address antifungal resistance.

## 1 Introduction

In an era of infectious disease outbreaks the ability to rapidly respond to emerging pathogens is critical and depends not only on the availability of molecular data but also the extrapolation of such data into clinical practice. Invasive fungal infections represent a growing global health burden, with increasing morbidity and mortality driven in part by the emergence of antifungal drug resistance. Such infections are an increasing threat to immunosuppressed patients and the elderly with growing number of incidences every year (1). Among clinically prevalent fungal pathogens, *Candida* and *Aspergillus* species account for the majority of infections (1). The over use of antifungals such as azoles and echinocandins has caused a shift to the epidemiology of pathogenic fungal species, leading to the emergence of antimicrobial drug resistance (AMR) (1, 2). The growing public health burden of AMR is well illustrated by the emergence of *Candidozyma auris* (*Candida auris)* which has been flagged by the US Centers for Disease Control and Prevention (CDC) as a serious public health threat (3, 4). The World Health Organization (WHO) has classified *C. auris* as a ‘critical priority’ pathogen in its fungal priority pathogens list (FPPL) underscoring urgent public health action and the European Union (EU) has launched coordinated initiatives to through a One Health approach to curb AMR (2, 5).

*C. auris* has caused serious outbreaks worldwide since its discovery in 2009, reported in over 40 countries (3, 4, 6). Genomic analysis of *C. auris* clinical strains have identified six clades, based on geographic location including clade I (South Asia), clade II (East Asia), clade III (Africa), clade IV (South America), clade V (Iran) and the very recent clade VI (Singapore), with clade I being the most prevalent (6–8). The pathogen has been associated with deep-seated infections of the bloodstream (candidemia), wound, respiratory tract, urinary tract, and ear infections, with high mortality rates ranging from 28% to 56% (3, 4). Furthermore, several outbreaks of *C. auris* have been reported worldwide since the emergence of coronavirus disease 2019 (COVID-19) (9), and co-infections with COVID-19 has resulted in a mortality rate of over 80% in some instances (9, 10).

The clinical challenges posed by *C. auris* stems from several sources including the fact that it is often misdiagnosed with other *Candida* species and requires rigorous molecular biology techniques for identification (11). Another bottleneck is the pathogens’ high transmissibility, where rapid patient to patient transmission has been reported (3, 4). In addition, *C. auris* possess a high degree of environmental persistence including tenacious contaminations of inanimate objects such as catheters and persistent colonization of the skin, thus making it difficult to control and eradicate outbreaks from affected areas (3, 4). For such reasons outbreaks can last for several months or even years before they are completely uprooted from hospitals.

A major impediment to combat *C. auris* is its multi-drug resistance (MDR) profile. It has been reported that 90% of *C. auris* clinical isolates are resistant to fluconazole, 35% to Amphotericin B (AmB), 7% to echinocandins, 3% to flucytosine and over 40% were reported to be resistant to more than 2 classes of antifungals (12). Treatment options therefore can be limited by intrinsic and secondary resistance and due to the availability of low number of antifungal families. This is particularly relevant in case of AmB resistance, which comes at a high cost because such a resistance essentially eliminates the last therapeutic option for treatment (13). Despite the effectiveness of the polyene AmB, its clinical use is limited in most countries due to its severe side-effects, such as nephro- and hepatotoxicity (14). The mode of action of this drug still remains enigmatic. The paradigm seems to be fixed at the pore formation theory, which states that AmB acts by forming pores on the cell membrane after binding to ergosterol thereby effectuating osmolysis (14). However, within this context, the fungicidal effect of AmB has been proposed to be more complex with induction of apoptosis and oxidative damage being implicated (14). To combat *C. auris* the above challenges need to be addressed in order to enable better infection control and preventive measures against the pathogen. Therefore, the quest for novel compounds with promising anti-*Auris* activity remains relentless. Compounds with potent anti-*Auris* can not only be used with a scope of potential therapy but can also be employed to limit the spread of the microorganism under clinical setting including in hospitals.

Drug repurposing (drug repositioning) is an attractive strategy to expand the application of existing approved drugs, considering the high attrition rates, extensive costs and time-consuming process for *de-novo* drug development (15). Even with a fast-tracked approval process of *de-novo* molecules as seen during the COVID-19 vaccine development, public mistrust often places such efforts back to time-consuming process and hinders our ability to combat pathogens. Repurposed drugs can be considered ‘de-risked’ compounds that have already passed safety assessment in preclinical models and in humans if early-stage trials have been completed, therefore are more likely to pass in subsequent efficacy trials.

Therefore, in order to expand the molecular knowledge spectrum of *C. auris* and address some of the key challenges associated with the pathogen, in this study, we applied a tailored drug repurposing strategy to identify FDA-approved compounds with activity against *C. auris*, coupled with quantitative proteomic analyses and electron microscopy, to elucidate potential antifungal resistance mechanisms employed by *C. auris* including those associated with AmB.

## 2 Materials and Method

### 2.1 *In-silico* drug screening strategy and generation custom drug library

The drug-repurposing strategy was guided by an *in-silico* drug screening approach based on Basic Local Alignment Search Tool (blast) (Figure 1) (16). In brief, drug target sequences were downloaded from DrugBank database *(version_2021)* consisting of protein targets of FDA approved drugs (17). The reference proteome sequences of *C. auris* were downloaded from the UniProt database and a blastp algorithm was carried out against the DrugBank target sequences (18, 19). Compounds identified with E-value: 0.00 were utilized to construct a custom drug library, from which at least one representative drug per unique *C. auris* protein ID was selected, yielding a targeted narrowed down set of 14 candidate compounds to be tested for primary *in-vitro* screening.

**Figure 1:**
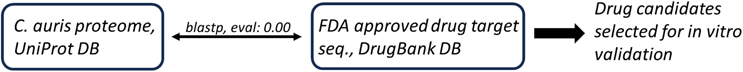
Drug repurposing strategy aided by using blastp screening procedure using *C. auris* protein sequences from UniProt database and DrugBank protein target sequences.

### 2.2 Candida strains and culture

In this study we used *C. auris* isolates belonging to five clades including AR389 (clade I), CBS10913 (clade II), AR383 (clade III), AR385 (clade IV) and AR1097 (clade V). Additionally, *Candida albicans* (SC5314) and *Candida glabrata* (ATCC2001) were also utilized. The candida cells were routinely cultured in YPD liquid media (1% yeast extract, 2% peptone, and 2% glucose) at 30°C with constant shaking at 200 rpm.

### 2.3 *In-vitro* screening assay

Primary screening of selected compounds was performed using spot assays to determine fungal growth inhibition. In brief, plates were prepared using media RPMI-1640 (Sigma-Aldrich, USA) buffered with 0.165mol/L 3-(N-morpholino) propanesulfonic acid (MOPS) (Sigma-Aldrich; USA), 2% glucose at pH 7.0 and 2% agar. Prior to experimentation, fungal cells were streaked onto YPD agar plates and allowed to propagate for 3 days, followed by cultures in YPD broth overnight at 30°C in 200 rpm shaker. Next fresh cultures were prepared and allowed to reach an optical density of 600 nm (OD_600_) corresponding to 1.0. Following which fungal cells were washed 3x in RPMI-MOPS media and adjusted to working inoculums of OD_600_ 0.2 cell suspension, from which 200μL was seeded into 96 well plates to be used by a robot for spotting. All the tested compounds were prepared in dimethyl sulfoxide (DMSO). Plates were created with a final drug concentration of 50μM (20) and 2μg/mL for AmB. The following compounds were selected from the *in-silico* screening library: Carboxin (mw: 235.30), Cladribine (mw: 285.69), Colchicene (mw: 399.43), Dasatinib (mw: 488.01), Enasidenib (mw: 473.38), Fostamatinib (mw: 580.46), Griseofulvin (mw: 352.77), Mupirocin (mw: 500.62), Pemetrexed (mw: 427.41), Pimecrolimus (mw: 810.45), Selinexor (mw: 443.31), Sulfinpyrazone (mw: 404.12), Tavaborole (mw: 151.93), Triclabendazole (mw: 359.65). After spotting by robotic arm, the plates were incubated at 30°C and growth inhibition was visually evaluated at 24h and 48h. Any compound displaying fungal growth inhibition across clades over 48h was monitored for up to 120h. Following which antifungal susceptibility test (AST) by broth dilution was carried out to determine minimum inhibitory concentration (MIC). In brief, AST was performed using media RPMI-1640 buffered with MOPS and 2% glucose at pH 7.0. Fungal cell suspensions of OD_600_ 0.2 were generated from which 100μL was seeded into 96-well plates, followed by the addition of 100μL 2-fold serial dilution series of positive hit compound. Controls wells contained no drug and experiments were performed as triplicates. The 96-well plates were then incubated at 30°C in 200 rpm shaker and growth inhibition was evaluated after 24h by measuring the OD_600_ using a Victor Nivo plate reader (PerkinElmer, USA).

### 2.4 Proteome isolation

In brief, *C. auris* clade I cells were grown to OD_600_ 0.8 in RPMI-MOPS with 2% glucose at 30°C in 200 rpm shaker followed by cell count by a CASY counter (Roche, Swiss). Following which, 5mL fresh cultures using the same media were seeded with 1 x 10^7^ candida cells for treatment with 2μM AmB or Tavaborole for 4hrs. All samples including untreated controls contained 0.002% DMSO. Experiments were performed as biological triplicates per treatment group. Following the treatment, protein was isolated from fungal cells. Briefly, candida cells were centrifuged at 3000 rpm for 5 min, followed by ice cold PBS wash-3x after which the cell pellet was resuspended in 500μL of ice-cold candida lysis buffer (1% sodium deoxycholate (SDC), 100mM Tris-HCl, 150mM NaCl, 1mM PMSF, 1mM EDTA, 1x cOmplete™ Protease Inhibitor (Roche, Swiss)) in 1.5mL screw-cap tubes (Sarstedt, Germany). Next, the candida cells were subjected to mechanical disruption by glass beads using FastPrep™-24 5G Bead Beating (Fisher Scientific, USA). Following which, a small hole was punctured at the bottom of the screw-cap tube using red-hot flamed needle and the screw-cap tube was then inserted into 1.5mL Eppendorf tube and centrifuge at 3000 rmp for 2 min at 4°C to separate the lysate from the glass beads. The lysate collected in the Eppendorf tube was then subjected to acetone precipitation by adding four volumes of ice-cold 100% acetone followed by 2h incubation at -20°C. Post incubation, the samples was centrifuged at 3000 rpm for 5min to obtain the resulting protein pellet and supernatant acetone was discarded. The protein pellet was air dried on ice and was further processed for mass spectrometric analysis.

### 2.5 Mass spectrometry sample preparation

Briefly, protein pellets were resuspended in 50µL 4%(w/v) SDS, 100mM Tris/HCl pH 8.5, 0.1M DTT, shaken at RT until fully resuspended and incubated at 95°C for 4 min. Lysates were clarified by centrifugation at 16000g for 10 min at 20°C. Supernatants were transferred to new tubes and protein concentration was measured using 600nm protein assay kit (Pierce) with SDS compatibility reagent. Solutions were diluted 1:10 with dH2O, transferred to FASP filters in two steps, and centrifuged for 20 min at 12000*g* each step. Filters were washed with 200µL 8M urea in 100mM Tris/HCl pH 8.5 for 20 min at 12000*g*, 100µL 50mM iodoacetamide in 100mM Tris/HCl pH 8.5 were added, shaken for 1 min and incubated 30 min in the dark at room temperature. Filters were centrifuged for 10 min at 12000g followed by washing three times with 200µL 8M urea in 100mM Tris/HCl pH 8.5 and three times with 100µL 100mM Tris/HCl pH 8.5. Filters were transferred to new collection tubes, 40µL 50mM ABC containing 1µg trypsin (Promega) were added and kept at 37°C overnight in a wet chamber to prevent evaporation. Digested peptides were collected by centrifuging for 20 min at 12000*g*. Filters were washed with 40µL 100mM Tris/HCl pH 8.5, centrifuged for 15 min at 12000*g* resulting in pooled eluates. Digested peptides were acidified with 10µL 10% TFA and the peptides were desalted using an MCX 96 well plate (Waters).

### 2.6 Liquid chromatography separation coupled to mass spectrometry

Peptides were separated on an Ultimate 3000 RSLC nano-flow chromatography system (Thermo-Fisher), using a pre-column for sample loading (Acclaim PepMap C18, 2 cm × 0.1 mm, 5μm, Thermo-Fisher), and a C18 analytical column (Acclaim PepMap C18, 50 cm × 0.75mm, 2μm, Thermo-Fisher), applying a segmented linear gradient from 2% to 35% and finally 80% solvent B (80 % acetonitrile, 0.1 % formic acid; solvent A 0.1 % formic acid) at a flow rate of 230nL/min over 120 min. Eluting peptides were analyzed on an Exploris 480 Orbitrap mass spectrometer (Thermo Fisher), which was coupled to the column with a FAIMS pro ion-source (Thermo-Fisher) using coated emitter tips (PepSep, MSWil).

### 2.7 Mass spectrometry data acquisition in data-independent acquisition mode (DIA) and raw data analysis

The mass spectrometer was operated in DIA mode with the FAIMS CV set to -45, the survey scans were obtained in a mass range of 350-1200 m/z, at a resolution of 30k at 200 m/z and a normalized AGC target at 300%. 31 MSMS spectra with variable isolation width between 14 and 27 m/z covering 399.5-899.5 m/z range including 1 m/z windows overlap, were acquired in the HCD cell at 30% collision energy at a normalized AGC target of 1000% and a resolution of 30k. The max. injection time was set to auto. Raw data were processed using Spectronaut software (version 16.1.220730.53000, https://biognosys.com/software/spectronaut/) with the DirectDIA workflow. The Uniprot *C. auris* reference proteome (version 2022_02, www.uniprot.org), as well as a database of most common contaminants were used. The searches were performed with full trypsin specificity and a maximum of 2 missed cleavages at a protein and peptide spectrum match false discovery rate of 1%. Carbamidomethylation of cysteine residues were set as fixed, oxidation of methionine and N-terminal acetylation as variable modifications. The global normalization and imputation were done in Spectronaut - all other parameters were left at default.

### 2.8 Data analysis using R scripts

Spectronaut output tables were further processed using Cassiopeia_LFQ 4.6.4 (https://github.com/maxperutzlabs-ms/Cassiopeia_LFQ). Contaminant proteins, protein groups identified only by one peptide and protein groups with less than two quantitative values in one experimental group, were removed for further analysis. Differences between groups were statistically evaluated using the LIMMA package (21) at 5% FDR (Benjamini-Hochberg).

### 2.9 Proteomics data deposition

The mass spectrometry proteomics data have been deposited to the ProteomeXchange Consortium via the PRIDE partner repository (22) with the dataset identifier PXD057542.

### 2.10 Transmission electron microscopy (TEM)

An aliquot of fungal cells used for proteomics was also subjected to electron microscopy. In brief, following the drug treatment, candida cells were pelleted at 3000 rpm for 5min. Cell pellets were then fixed using a mixture of 2% glutaraldehyde (Agar Scientific, UK) and 2% paraformaldehyde (Electron Microscopy Sciences, USA) in 0.1mol/l sodium cacodylate buffer, pH 7.2 at room temperature overnight followed by 3 rinsing steps with the same and a post-fixation in 2% osmium tetroxide (Agar Scientific, UK) in 0.1mol/l sodium cacodylate buffer. Dehydration was performed in a graded series of acetone and samples were embedded in Agar 100 resin (Agar Scientific, UK). 70-nm sections were cut and post-stained with 2% uranyl acetate and Reynolds lead citrate (Delta Microscopies, France). Micrographs were recorded on an FEI Morgagni 268D (FEI, Netherlands) operated at 80 keV, equipped with a Mega View III CCD camera (Olympus-SIS).

## 3 Results

### 3.1 *In-silico* drug screening

Following the *in-silico* drug screening procedure up to 43 unique compounds were identified targeting 24 *C. auris* protein targets (Supplementary Table 1). Established frontline antifungals such as the Azole group (Fluconazole, Oxiconazole, Terconazole, Clotrimazole, Voriconazole, Tioconazole, Miconazole, Econazole, Sertaconazole, Posaconazole, Bifonazole, Luliconazole), Echinocandin class (Anidulafungin, Caspofungin, Micafungin) and Flucytosine (5-FC) were identified via the *in-silico* screening strategy, serving as an important reference benchmark to validate the procedure. The narrowed down custom list for the primary *in-vitro* screen targeting at least one *C. auris* protein included 14 compounds such as Carboxin, Cladribine, Colchicene, Dasatinib, Enasidenib, Fostamatinib, Griseofulvin, Mupirocin, Pemetrexed, Pimecrolimus, Selinexor, Sulfinpyrazone, Tavaborole and Triclabendazole.

### 3.2 *In-vitro* primary drug screening

Primary *in-vitro* spot assays performed on the 14 repurposed compounds identified two positive hits as novel indication of an existing drug; Triclabendazole and Tavaborole, with the latter emerging as the lead compound exhibiting sustained inhibition across *C. auris* clades beyond 48h. The compound Tavaborole (Kerydin/AN2690) (mw: 151.93) (Figure 2) displayed robust effect against all five tested clades of *C. auris,* plus on *C. albicans and C. glabrata* for up to 120h; a threshold time-point to determine fungicidal vs fungistatic effect (Figure 3). Drug susceptibility test confirmed Tavaborole activity against *Candida* species with a low MIC range of 3µM (Figure 4). The anthelmintic drug Triclabendazole exhibited modest anti-*Candida* activity (Supplementary Material 1), however, its effect was weaker than that of Tavaborole. Additionally, a recent study (23), has already identified and confirmed the antifungal activity of Triclabendazole against *C. auris* clades, as such it was not investigated further and was beyond the scope of this current study.

**Figure 2:**
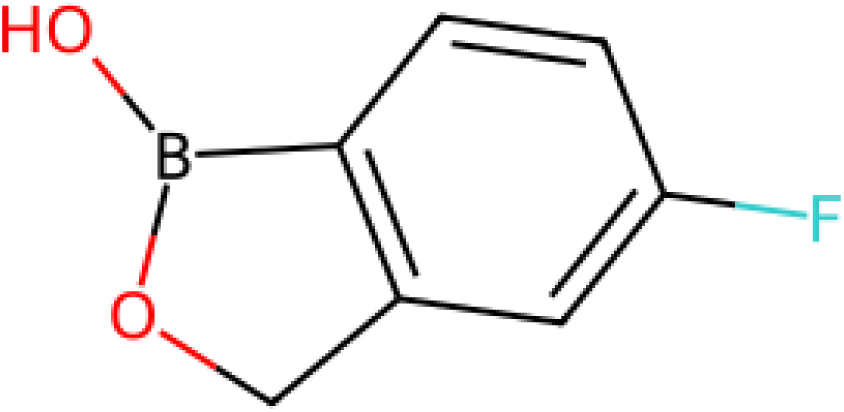
Chemical structure of Tavaborole also sold under the brand name Kerydin (AN2690). Systematic name: 5-fluoro-1,3-dihydro-1-hydroxy-2,1-benzoxaborole, chemical formula C7H6BFO2. Diagram generated using open-source cheminformatics tool RDKit.

**Figure 3:**
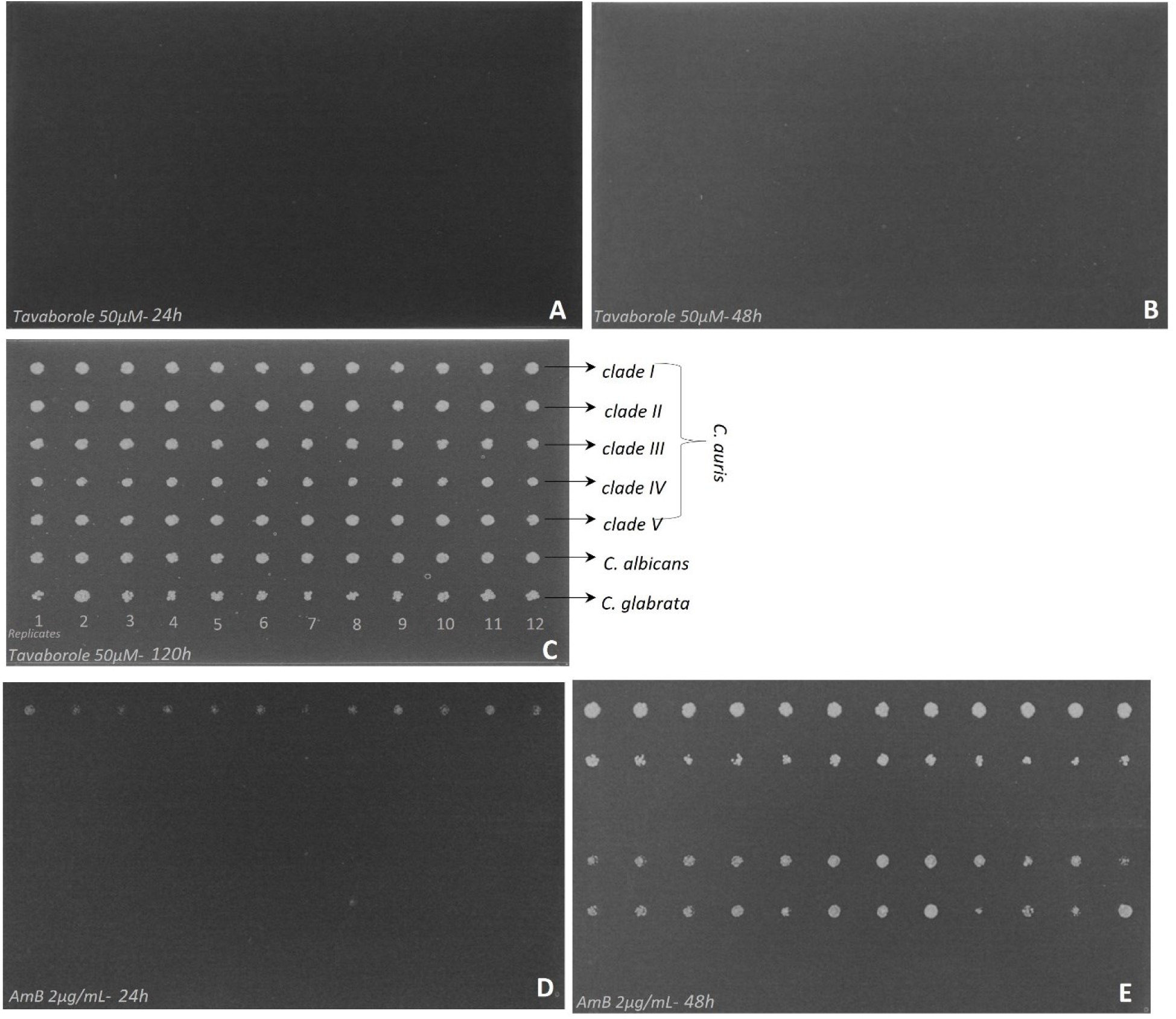
Comparison of the primary *in-vitro* screening of Tavaborole (50 µM) and AmB (2µg/mL) against *C. auris* clades I–V, *C. albicans* and *C. glabrata*. (A-C) No visible fungal growth was observed in the presence of Tavaborole for up to 120h (fungistatic) of incubation. (D-E) Limited fungal growth was observed following 48h of incubation with AmB.

**Figure 4:**
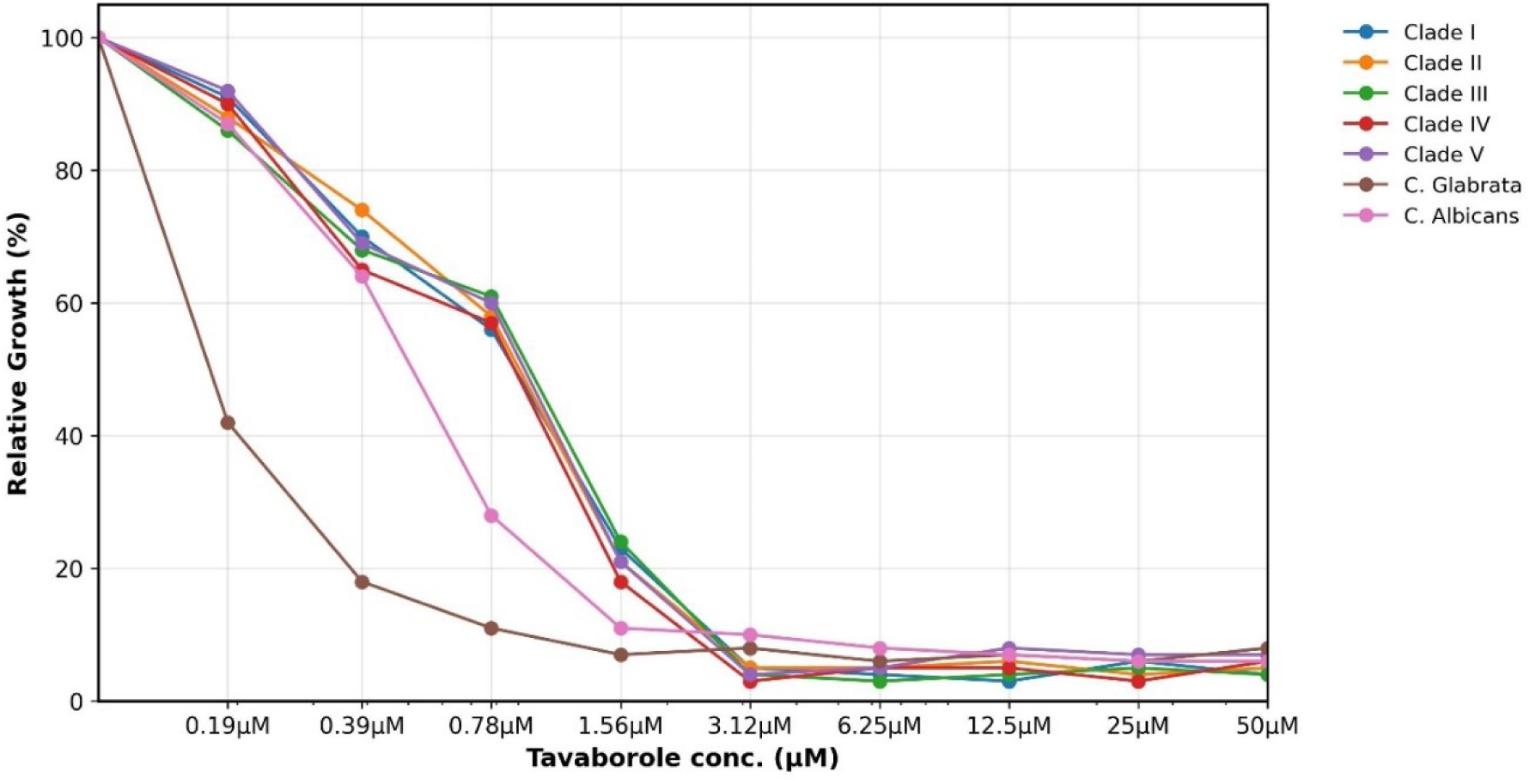
Antifungal susceptibility test of *C. auris* isolates from clades I-V, *C. albicans* and *C. glabrata* against Tavaborole using broth microdilution assay, MIC determined at lowest concentration achieving ≥90% fungal growth inhibition.

### 3.3 Proteomics

Proteomics analysis identified 4555 *C. auris* proteins which covers 84% of the *C. auris* UniProt proteome database of 5409 sequences. The treatment with Tavaborole caused the differential regulation of 151 proteins (Supplementary Table 1) with the increased abundance of proteins including LEU1, LEU4, orf19.1502, GLT1, GCN4, HPA2, orf19.813, SOD1, YHB1 and GLG21 (Figure 5A). Upon AmB treatment 50 proteins were differentially regulated (Supplementary Table 1) with the increased abundance of proteins including DDR48, orf19.7085, orf19.813, RHR2, LEU1, and GLG21 (Figure 5B).

**Figure 5:**
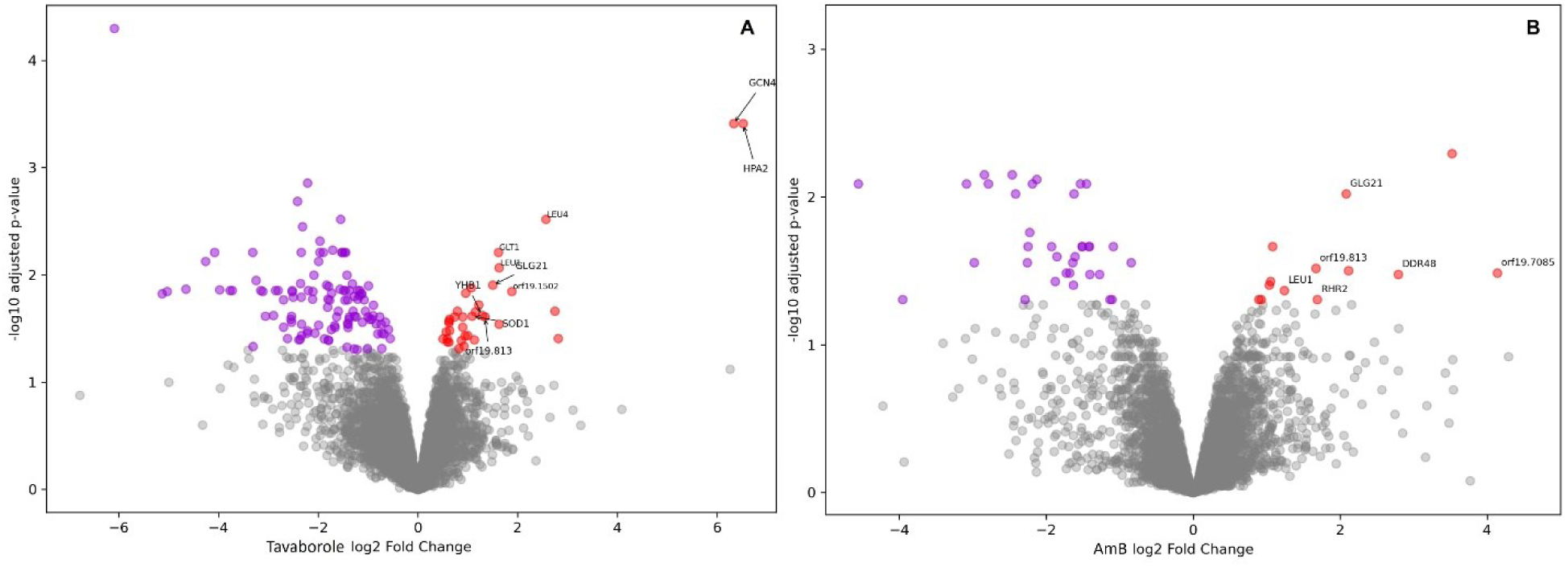
Visualization of quantitative proteomics data from *C. auris* following antifungal treatment. Volcano plots depict differentially abundant proteins upon exposure to (A) Tavaborole and (B) polyene AmB. Significantly regulated proteins were defined using an adjusted P value ≤ 0.05.

### 3.4 TEM imaging analysis

Electron microscopy was performed on *C. auris* cells treated with Tavaborole and AmB (Figure 6). Untreated candida cells displayed well defined cell-wall and plasma membrane with homogeneous cytoplasm and intact intracellular organization. Candida treated with Tavaborole exhibited pronounced intracellular alterations characterized by enlarged vacuoles and compromised internal architecture, suggesting intracellular stress and potential perturbation cellular homeostasis, while the cell-wall and plasma membrane remained largely intact. In contrast, candida cells treated with AmB showed structural damage characterized by extensive vacuolization, separation between plasma membrane and cytoplasmic contents, and signs of compromised membrane integrity.

**Figure 6:**
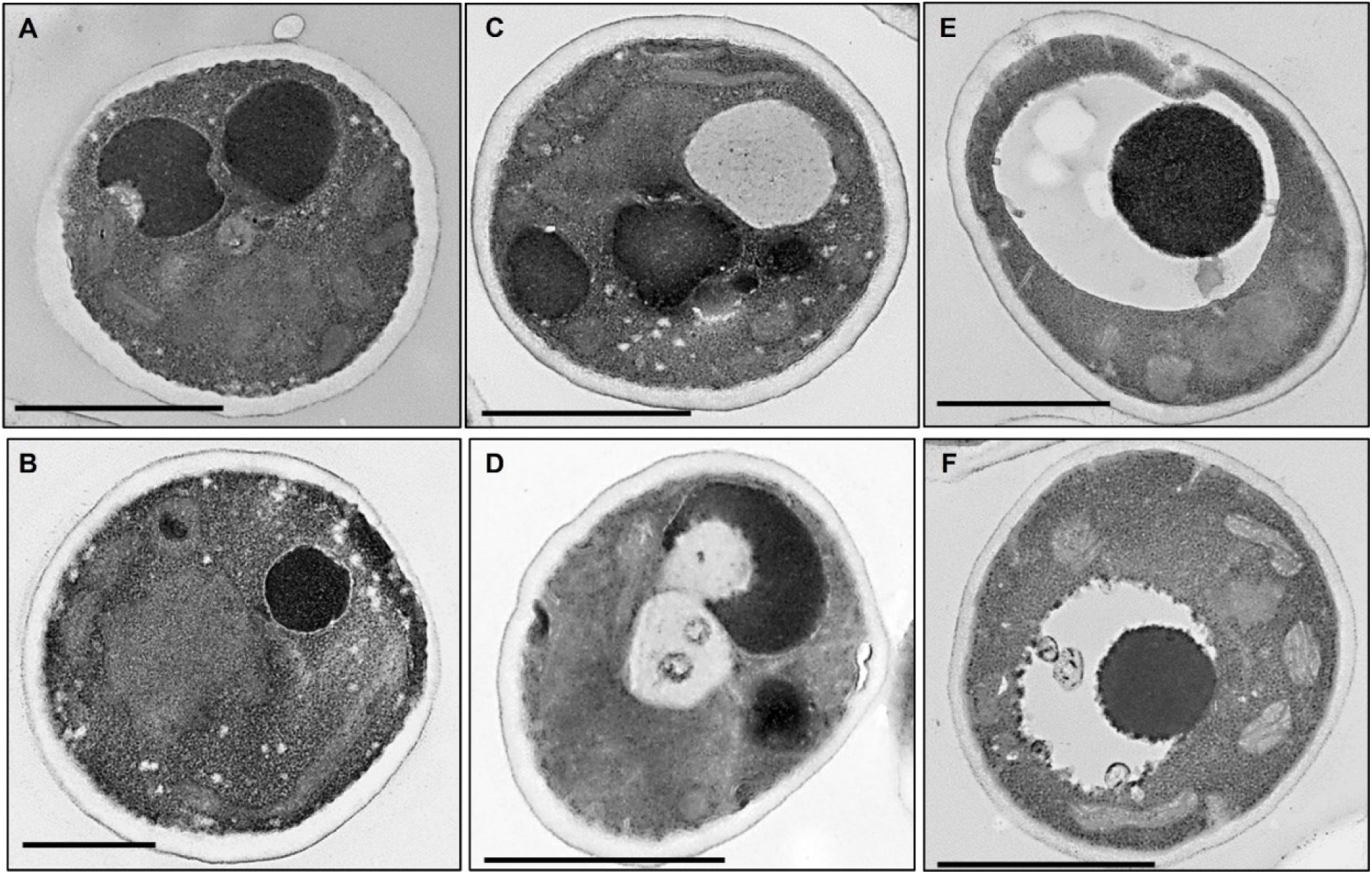
Transmission electron microscopy (TEM) of *C. auris* cells treated with Tavaborole and AmB; (A-B) untreated control *C. auris*, (C-D) candida cells treated with Tavaborole and, (E-F) *C. auris* treated with AmB. Scale bar = 1 μm.

## 4 Discussion

Infections caused by *C. auris* pose a major challenge in regards to its treatment, particularly due to its drug resistance profile (24). Importantly, resistance to the polyene AmB is of particular concern as it is rare among fungal pathogens including *Candida* species (25). Resistance to AmB essentially eliminates the last therapeutic option against fungal pathogens (13). The mechanism of AmB action remains enigmatic, though reports suggest that the drug interacts with fungal cell membrane ergosterol thereby effectuating osmolysis (14). Notably, AmB resistance has been reported across *C. auris* clades and pose a massive health threat across the globe. As such, a deeper understanding of *C. auris* drug resistance mechanism as well as the quest for novel antifungal alternatives is of immense importance to combat this enigmatic pathogen.

In an era of precision medicine and artificial intelligence (AI) driven drug discovery, the demand for molecules with a high likelihood of therapeutic efficacy is of monumental value. To this end, we applied a tailored drug repurposing strategy to generate a custom drug screening library consisting of 14 compounds, predicted to display anti-*Candida* activity with high probability. Our approach was guided by utilizing the *C. auris* proteome sequences to perform a blast screen against FDA approved drug target sequences from DrugBank database. With this procedure we were able to dramatically reduce the number of drugs to be tested from thousands to only a handful. In general, antifungal drug screening libraries are expensive and consist of over thousands of compounds to be tested, and the screening usually result in a few successful ‘hit’ compounds (26). For example, the Johns Hopkins Clinical Compound Library consisted of over 1500 compounds to be tested with six successful hits, the Prestwick library comprised of 1200 compounds with seven hits and the TargetMol library of FDA-approved drugs contained 1068 compounds for screening with one successful hit (26, 27). In contrast, our drug screening library consisted of 14 compounds to be tested leading to two successful hits, therefore the ratio of the number of drugs tested to yield one successful hit is relatively high. Large-scale repurposing screens typically yield hit rates below 1%, including the Johns Hopkins Clinical Compound Library (0.40%), Prestwick Chemical Library (0.58%), and the TargetMol FDA-approved drug library (0.094%), whereas our targeted screening strategy identified 2 hits from 14 compounds (14.29%), representing an approximately 25 to 150-fold enrichment in hit rate. The drug Tavaborole identified as the lead hit, displayed potent fungistatic activity against clinical isolates of five *C. auris* clades, and *C. albicans* and *C. glabrata*. The second identified drug Triclabendazole also displayed antifungal activity against *C. auris* corroborating with the recent study (23).

Tavaborole is an FDA approved drug is sold under the name Kerydin, which is used to treat fungal infection onychomycosis caused by *Trichophyton rubrum* and *Trichophyton mentagrophytes*. The drug exerts its antifungal activity by inhibiting Leucyl-tRNA synthetase (LeuRS) to disrupt fungal protein synthesis (28). Proteomics analysis revealed that upon exposure of *C. auris* cells to Tavaborole several proteins were over abundant namely LEU1, LEU4, orf19.1502, GLT1, GCN4, HPA2, orf19.813, SOD1, YHB1 and GLG21. Tavaborole inhibits Leucyl-tRNA synthetase which catalyzes ligation of the amino acid L-leucine to tRNA (29). Here, the exposure of *C. auris* to Tavaborole induced the upregulation of leucine biosynthesis associated protein LEU1 and LEU4 (30). Furthermore, the orf19.1502 associated protein which has a predicted aminoacyl-tRNA hydrolase activity was upregulated (31). An aminoacyl-tRNA is an tRNA with its cognate amino acid. Every amino acid has its own specific aminoacyl-tRNA synthetase which chemically binds it to tRNA in order to be transferred to a growing peptide (32). Under certain circumstances where a wrong amino acid forms a cognate tRNA, it must be hydrolyzed in order to prevent incorrect protein synthesis (32, 33). The upregulation of the above *C. auris* proteins suggests an intricate control of amino acid biosynthesis in response to Tavaborole. This is further consolidated by the display of general amino acid control, also known as the GCN response, a phenomenon where deprivation of a particular amino acid induces the expression of genes of all amino acid biosynthesis pathways to increase (34). This is evident in our study by the positive regulation of GLT1 responsible for glutamate biosynthesis (35). In *C. albicans* GCN4 functions as a transcriptional regulator of amino acid biosynthesis including coordinating responses to amino acid starvation (34). The transcription modulator GCN4 can employ histone acetyltransferase (HATs) complexes to confer transcriptional activation (36). The HPA2 is a member of HATs that can modify histones by acetylating lysine residues at histone tails or at histone globular domains (37–39). Whereby, histone modifications can regulate the transcriptional state of genes and can confer various advantages including responding to external stimuli (37–40). For *C. albicans* it has been reported that HATs regulate genes to respond to external stimuli in the context of virulence, oxidative stress and antifungal drug tolerance (41). Additionally, the upregulation of small heat shock protein associated with orf19.813, superoxide dismutase SOD1 and nitric oxide dioxygenase YHB1 indicates a possible oxidative stress response (42–44). Furthermore, the role of glycogen metabolism was highlighted with the upregulation of GLG21, a homolog of *S. cerevisiae* GLG2 encoding the enzyme glucosyltransferase mediating glycogen metabolism (31, 45). Glycogen metabolism in candida has been linked to virulence, survival under stress environment and synthesis of cell-wall (45–47). The upregulation of GLG21 upon Tavaborole exposure thus seem to be a protective mechanism employed by *C. auris* to circumvent potential stress induced by the drug.

The proteome dynamics of *C. auris* upon exposure to AmB revealed the upregulation of several proteins associated with DDR48, orf19.7085, orf19.813, RHR2, LEU1 and GLG21. The polyene caused the increased abundance of stress response molecules including proteins associated with DDR48, orf19.7085 and orf19.813. Stress adaptation is a crucial factor for microbial survival under dynamic environments including candida species. Several studies have suggested that AmB can autoxidize to bring about oxidative stress with a fungicidal impact (25). In *C. albicans* it has been reported that AmB induces oxidative and nitrosative stress by production of reactive oxygen species (ROS) and nitrogen species (48–50). The stress protein DDR48 was reported to be induced in *C. albicans* in response to oxidative stress, antifungal drugs, exposure to cell wall-perturbing agents and DNA damage (51). The proteins associated with orf19.7085 and orf19.813 are also classified as oxidative stress molecules (31). Therefore, it can be inferred, that the above stress response molecules expressed by *C. auris,* underpins the effect of oxidative stress induced by AmB and reflects on the pathogens’ stress adaptation mechanism including antifungal resistance. The RHR2 which encodes for glycerol-3-phosphatase involved in glycerol biosynthesis and has been reported to be mediating fungal stress response including osmotic stress and oxidative stress (52, 53). Increased osmolarity is known to cause water loss and cell shrinking, therefore a major survival strategy is to accumulate suitable osmolytes such as glycerol to maintain water balance and restore cell volume (54). In yeast osmotic stress causes overproduction of glycerol which in turn is mediated by the high-osmolarity glycerol (HOG) mitogen-activated protein kinase (MAPK) pathway (55, 56). This osmosensing pathway has been implicated in pathogenicity and cell wall biogenesis in *C. albicans* (56). Previous proteomics study in *C. albicans* reported the overabundance of RHR2 protein upon exposure to AmB which could be indicative of glycerol buildup (55). Given that AmB functions by creating pores on fungal cell wall to effectuate osmolysis, the overabundance of RHR2 in our study suggests that *C. auris* could employ excess glycerol as an osmolyte to counter the osmotic stress caused by AmB. Finally, the common upregulation of GLG21, LEU1 and orf19.813 associated proteins upon exposure to both Tavaborole and AmB suggests, an intrinsic drug resistance response in *C. auris,* in which the organism adapts its glycogen metabolism and amino acid biosynthesis coupled with stress responses to counter antifungal agents. Furthermore, electron microscopic analysis of *C. auris* upon exposure to Tavaborole corroborates with the suggestive mode of action of the drug, indicating intracellular stress inducement and potential disruption of cellular homeostasis. In contrast, exposure to AmB is indicative of a more profound adverse effect on candida cells suggesting severe cellular stress and compromising membrane integrity, consistent with the polyene’s hypothesized mode of action of inducing oxidative stress and membrane disruption.

## 5 Conclusion

In this study, we applied a custom drug-repurposing strategy to identify novel indications for existing compounds, leading to the identification of Tavaborole as a potent fungistatic agent against *C. auris*. Compared to commercial drug screen libraries, our targeted drug-repurposing procedure resulted in a much higher hit rate, which can be readily adapted to target other organisms and enhanced using AI tools. The lead compound Tavaborole exhibited robust anti-*Candida* activity across the five tested clades of *C. auris*, as well as *C. albicans* and *C. glabrata*. Additionally, to elucidate antifungal resistance mechanisms in *C. auris*, we performed quantitative proteomic analyses following fungal exposure to Tavaborole and AmB, complemented by electron microscopy to assess drug-induced ultrastructural changes. Our analyses indicated that *C. auris* mounts distinct yet overlapping adaptive responses to antifungal stress. Tavaborole exposure primarily induced stress response pathways and modulation of amino acid biosynthesis. In contrast, AmB triggered a broader, multi-layered resistance response involving oxidative stress adaptation, osmolyte production and metabolic reprogramming encompassing glycogen metabolism and amino acid biosynthesis. Notably, alterations in glycogen metabolism and amino acid biosynthesis were common to both Tavaborole and AmB treatments, suggesting conserved antifungal adaptation mechanisms that warrant further investigations.

## 6.#Acknowledgment

This study was funded by the Austrian Science Fund (FWF) project CandidOmics-P33425 (Grant DOI: 10.55776/P33425). Proteomics analyses were performed by the Mass Spectrometry Facility at Max Perutz Labs using the VBCF instrument pool. Electron microscopy imaging was done by the Electron Microscopy Facility, Vienna BioCenter Core Facilities (VBCF), Vienna, Austria. We would like to thank Karl Kuchler and lab members for their support.

## 7 Contributions

Conceptualization: RM. Methodology: RM. Software: RM. Validation: RM, AB. Formal analysis: RM. Investigation: RM, AB. Resources: RM. Data curation: RM. Visualization: RM. Supervision: RM. Project administration: RM. Funding acquisition: RM. Writing-original draft: RM. Writing-Review & Editing: Both authors.

## 8 Conflict of interest

The authors declare no competing interests.

## Notes

### Competing Interest Statement

The authors have declared no competing interest.

## References

1. Vitiello A, Ferrara F, Boccellino M, Ponzo A, Cimmino C, Comberiati E, Zovi A, Clemente S, Sabbatucci M. 2023. Antifungal Drug Resistance: An Emergent Health Threat. Biomedicines 11:1063.

2. Fisher MC, Burnett F, Chandler C, Gow NAR, Gurr S, Hart A, Holmes A, May RC, Quinn J, Soliman T, Talbot NJ, West HM, West JS, White PL, Bromley M, Armstrong-James D. 2024. A one health roadmap towards understanding and mitigating emerging Fungal Antimicrobial Resistance: fAMR. npj Antimicrobials and Resistance 2:36.

3. Nett JE. 2019. Candida auris: An emerging pathogen “incognito”? PLoS Pathog 15:e1007638.

4. Sears D, Schwartz BS. 2017. Candida auris: An emerging multidrug-resistant pathogen. International Journal of Infectious Diseases 63:95–98.

5. Davido B, Ny S, van Lingen C, Årdal C, Alonso Irujo L, Linnros S, et al. 2026. Strengthening antimicrobial resistance governance in Europe: a coordinated one health approach. The Lancet Regional Health - Europe 61:101540.

6. Hayes JF. 2024. Candida auris: Epidemiology Update and a Review of Strategies to Prevent Spread. J Clin Med 13:6675.

7. The Lancet Microbe. 2024. Candida auris: new clade, same challenges. Lancet Microbe 5:100977.

8. De Gaetano S, Midiri A, Mancuso G, Avola MG, Biondo C. 2024. Candida auris Outbreaks: Current Status and Future Perspectives. Microorganisms 12:927.

9. Tsai C-S, Lee SS-J, Chen W-C, Tseng C-H, Lee N-Y, Chen P-L, Li M-C, Syue L-S, Lo C-L, Ko W-C, Hung Y-P. 2023. COVID-19-associated candidiasis and the emerging concern of Candida auris infections. Journal of Microbiology, Immunology and Infection 56:672–679.

10. Villanueva-Lozano H, Treviño-Rangel R de J, González GM, Ramírez-Elizondo MT, Lara-Medrano R, Aleman-Bocanegra MC, Guajardo-Lara CE, Gaona-Chávez N, Castilleja-Leal F, Torre-Amione G, Martínez-Reséndez MF. 2021. Outbreak of Candida auris infection in a COVID-19 hospital in Mexico. Clinical Microbiology and Infection 27:813–816.

11. Lockhart SR, Lyman MM, Sexton DJ. 2022. Tools for Detecting a “Superbug”: Updates on Candida auris Testing. J Clin Microbiol 60.

12. Lockhart SR, Etienne KA, Vallabhaneni S, Farooqi J, Chowdhary A, Govender NP, Colombo AL, Calvo B, Cuomo CA, Desjardins CA, Berkow EL, Castanheira M, Magobo RE, Jabeen K, Asghar RJ, Meis JF, Jackson B, Chiller T, Litvintseva AP. 2017. Simultaneous Emergence of Multidrug-Resistant *Candida auris* on 3 Continents Confirmed by Whole-Genome Sequencing and Epidemiological Analyses. Clinical Infectious Diseases 64:134–140.

13. Vincent BM, Lancaster AK, Scherz-Shouval R, Whitesell L, Lindquist S. 2013. Fitness Trade-offs Restrict the Evolution of Resistance to Amphotericin B. PLoS Biol 11.

14. Mesa-Arango AC, Scorzoni L, Zaragoza O. 2012. It only takes one to do many jobs: Amphotericin B as antifungal and immunomodulatory drug. Front Microbiol 3:1–10.

15. Pushpakom S, Iorio F, Eyers PA, Escott KJ, Hopper S, Wells A, Doig A, Guilliams T, Latimer J, McNamee C, Norris A, Sanseau P, Cavalla D, Pirmohamed M. 2018. Drug repurposing: Progress, challenges and recommendations. Nat Rev Drug Discov. Nature Publishing Group 10.1038/nrd.2018.168.

16. Mazumdar R, Endler L, Monoyios A, Hess M, Bilic I. 2017. Establishment of a de novo Reference Transcriptome of Histomonas meleagridis Reveals Basic Insights About Biological Functions and Potential Pathogenic Mechanisms of the Parasite. Protist 168:663–685.

17. Wishart DS, Feunang YD, Guo AC, Lo EJ, Marcu A, Grant JR, Sajed T, Johnson D, Li C, Sayeeda Z, Assempour N, Iynkkaran I, Liu Y, Maciejewski A, Gale N, Wilson A, Chin L, Cummings R, Le D, Pon A, Knox C, Wilson M. 2018. DrugBank 5.0: a major update to the DrugBank database for 2018. Nucleic Acids Res 46:D1074–D1082.

18. Bateman A, Martin M-J, Orchard S, Magrane M, Ahmad S, Alpi E, Bowler-Barnett EH, et al. 2023. UniProt: the Universal Protein Knowledgebase in 2023. Nucleic Acids Res 51:D523–D531.

19. Knox C, Wilson M, Klinger CM, Franklin M, Oler E, Wilson A, Pon A, Cox J, Chin NE (Lucy), Strawbridge SA, Garcia-Patino M, Kruger R, Sivakumaran A, Sanford S, Doshi R, Khetarpal N, Fatokun O, Doucet D, Zubkowski A, Rayat DY, Jackson H, Harford K, Anjum A, Zakir M, Wang F, Tian S, Lee B, Liigand J, Peters H, Wang RQ (Rachel), Nguyen T, So D, Sharp M, da Silva R, Gabriel C, Scantlebury J, Jasinski M, Ackerman D, Jewison T, Sajed T, Gautam V, Wishart DS. 2024. DrugBank 6.0: the DrugBank Knowledgebase for 2024. Nucleic Acids Res 52:D1265–D1275.

20. Selin C, Stietz MS, Blanchard JE, Gehrke SS, Bernard S, Hall DG, Brown ED, Cardona ST. 2015. A Pipeline for Screening Small Molecules with Growth Inhibitory Activity against Burkholderia cenocepacia. PLoS One 10:e0128587.

21. Ritchie ME, Phipson B, Wu D, Hu Y, Law CW, Shi W, Smyth GK. 2015. limma powers differential expression analyses for RNA-sequencing and microarray studies. Nucleic Acids Res 43:e47–e47.

22. Perez-Riverol Y, Bai J, Bandla C, García-Seisdedos D, Hewapathirana S, Kamatchinathan S, Kundu DJ, Prakash A, Frericks-Zipper A, Eisenacher M, Walzer M, Wang S, Brazma A, Vizcaíno JA. 2022. The PRIDE database resources in 2022: a hub for mass spectrometry-based proteomics evidences. Nucleic Acids Res 50:D543–D552.

23. Lohse MB, Laurie MT, Levan S, Ziv N, Ennis CL, Nobile CJ, DeRisi J, Johnson AD. 2023. Broad susceptibility of *Candida auris* strains to 8-hydroxyquinolines and mechanisms of resistance. mBio 10.1128/mbio.01376-23.

24. Sharma C, Kadosh D. 2023. Perspective on the origin, resistance, and spread of the emerging human fungal pathogen Candida auris. PLoS Pathog 19:e1011190.

25. Mesa-Arango AC, Scorzoni L, Zaragoza O. 2012. It only takes one to do many jobs: Amphotericin B as antifungal and immunomodulatory drug. Front Microbiol 3.

26. Wall G, Lopez-Ribot JL. 2020. Screening Repurposing Libraries for Identification of Drugs with Novel Antifungal Activity. Antimicrob Agents Chemother 64.

27. Mei Y, Jiang T, Zou Y, Wang Y, Zhou J, Li J, Liu L, Tan J, Wei L, Li J, Dai H, Peng Y, Zhang L, Lopez-Ribot JL, Shapiro RS, Chen C, Liu N-N, Wang H. 2020. FDA Approved Drug Library Screening Identifies Robenidine as a Repositionable Antifungal. Front Microbiol 11.

28. He Z, Huang D-C, Guo D, Deng F, Sha Q, Zhang M-Z, Zhang W-H, Gu Y-C. 2023. Synthesis, fungicidal activity and molecular docking studies of tavaborole derivatives. Advanced Agrochem 2:185–195.

29. Mazzantini D, Celandroni F, Calvigioni M, Lupetti A, Ghelardi E. 2021. *In Vitro* Resistance and Evolution of Resistance to Tavaborole in Trichophyton rubrum. Antimicrob Agents Chemother 65.

30. Kohlhaw GB. 2003. Leucine Biosynthesis in Fungi: Entering Metabolism through the Back Door. Microbiology and Molecular Biology Reviews 67:1–15.

31. Skrzypek MS, Binkley J, Binkley G, Miyasato SR, Simison M, Sherlock G. 2017. The *Candida* Genome Database (CGD): incorporation of Assembly 22, systematic identifiers and visualization of high throughput sequencing data. Nucleic Acids Res 45:D592–D596.

32. Francklyn CS, Mullen P. 2019. Progress and challenges in aminoacyl-tRNA synthetase-based therapeutics. Journal of Biological Chemistry 294:5365–5385.

33. Jost J-P, Bock RM. 1969. Enzymatic Hydrolysis of N-Substituted Aminoacyl Transfer Ribonucleic Acid in Yeast. Journal of Biological Chemistry 244:5866–5873.

34. Tripathi G. 2002. Gcn4 co-ordinates morphogenetic and metabolic responses to amino acid starvation in Candida albicans. EMBO J 21:5448–5456.

35. Valenzuela L, Ballario P, Aranda C, Filetici P, González A. 1998. Regulation of Expression of *GLT1*, the Gene Encoding Glutamate Synthase in *Saccharomyces cerevisiae*. J Bacteriol 180:3533–3540.

36. Kuo M-H, vom Baur E, Struhl K, Allis CD. 2000. Gcn4 Activator Targets Gcn5 Histone Acetyltransferase to Specific Promoters Independently of Transcription. Mol Cell 6:1309–1320.

37. Conte M, Eletto D, Pannetta M, Petrone AM, Monti MC, Cassiano C, Giurato G, Rizzo F, Tessarz P, Petrella A, Tosco A, Porta A. 2022. Effects of Hst3p inhibition in Candida albicans: a genome-wide H3K56 acetylation analysis. Front Cell Infect Microbiol 12.

38. Jaiswal D, Turniansky R, Green EM. 2017. Choose Your Own Adventure: The Role of Histone Modifications in Yeast Cell Fate. J Mol Biol 429:1946–1957.

39. Kuchler K, Jenull S, Shivarathri R, Chauhan N. 2016. Fungal KATs/KDACs: A New Highway to Better Antifungal Drugs? PLoS Pathog 12:e1005938.

40. Lopes da Rosa J, Kaufman PD. 2012. Chromatin-mediated Candida albicans virulence. Biochimica et Biophysica Acta (BBA) - Gene Regulatory Mechanisms 1819:349–355.

41. Tscherner M, Stappler E, Hnisz D, Kuchler K. 2012. The histone acetyltransferase Hat1 facilitates DNA damage repair and morphogenesis in Candida albicans. Mol Microbiol 86:1197–1214.

42. Gong Y, Li T, Yu C, Sun S. 2017. Candida albicans Heat Shock Proteins and Hsps-Associated Signaling Pathways as Potential Antifungal Targets. Front Cell Infect Microbiol 7.

43. Martchenko M, Alarco A-M, Harcus D, Whiteway M. 2004. Superoxide Dismutases in *Candida albicans* : Transcriptional Regulation and Functional Characterization of the Hyphal-induced *SOD5* Gene. Mol Biol Cell 15:456–467.

44. Ullmann BD, Myers H, Chiranand W, Lazzell AL, Zhao Q, Vega LA, Lopez-Ribot JL, Gardner PR, Gustin MC. 2004. Inducible Defense Mechanism against Nitric Oxide in *Candida albicans*. Eukaryot Cell 3:715–723.

45. Miao J, Regan J, Cai C, Palmer GE, Williams DL, Kruppa MD, Peters BM. 2023. Glycogen Metabolism in Candida albicans Impacts Fitness and Virulence during Vulvovaginal and Invasive Candidiasis. mBio 14.

46. Loza L, Doering TL. 2024. A fungal protein organizes both glycogen and cell wall glucans. Proceedings of the National Academy of Sciences 121.

47. Larcombe DE, Bohovych IM, Pradhan A, Ma Q, Hickey E, Leaves I, Cameron G, Avelar GM, de Assis LJ, Childers DS, Bain JM, Lagree K, Mitchell AP, Netea MG, Erwig LP, Gow NAR, Brown AJP. 2023. Glucose-enhanced oxidative stress resistance—A protective anticipatory response that enhances the fitness of Candida albicans during systemic infection. PLoS Pathog 19:e1011505.

48. Sokol-Anderson ML, Brajtburg J, Medoff G. 1986. Amphotericin B-Induced Oxidative Damage and Killing of Candida albicans. Journal of Infectious Diseases 154:76–83.

49. Vriens K, Kumar PT, Struyfs C, Cools TL, Spincemaille P, Kokalj T, Sampaio-Marques B, Ludovico P, Lammertyn J, Cammue BPA, Thevissen K. 2017. Increasing the Fungicidal Action of Amphotericin B by Inhibiting the Nitric Oxide-Dependent Tolerance Pathway. Oxid Med Cell Longev 2017:1–17.

50. Mesa-Arango AC, Trevijano-Contador N, Román E, Sánchez-Fresneda R, Casas C, Herrero E, Argüelles JC, Pla J, Cuenca-Estrella M, Zaragoza O. 2014. The Production of Reactive Oxygen Species Is a Universal Action Mechanism of Amphotericin B against Pathogenic Yeasts and Contributes to the Fungicidal Effect of This Drug. Antimicrob Agents Chemother 58:6627–6638.

51. Cleary IA, MacGregor NB, Saville SP, Thomas DP. 2012. Investigating the Function of Ddr48p in Candida albicans. Eukaryot Cell 11:718–724.

52. Fan J, Whiteway M, Shen S-H. 2005. Disruption of a gene encoding glycerol 3-phosphatase from *Candida albicans* impairs intracellular glycerol accumulation-mediated salt-tolerance. FEMS Microbiol Lett 245:107–116.

53. Smith DA, Nicholls S, Morgan BA, Brown AJP, Quinn J. 2004. A Conserved Stress-activated Protein Kinase Regulates a Core Stress Response in the Human Pathogen *Candida albicans*. Mol Biol Cell 15:4179–4190.

54. de Nadal E, Posas F. 2022. The HOG pathway and the regulation of osmoadaptive responses in yeast. FEMS Yeast Res 22.

55. Hoehamer CF, Cummings ED, Hilliard GM, Rogers PD. 2010. Changes in the Proteome of *Candida albicans* in Response to Azole, Polyene, and Echinocandin Antifungal Agents. Antimicrob Agents Chemother 54:1655–1664.

56. Román E, Correia I, Prieto D, Alonso R, Pla J. 2020. The HOG MAPK pathway in Candida albicans: more than an osmosensing pathway. International Microbiology 23:23–29.

