## Supplementary Material 1 for "A targeted drug-repurposing strategy identifies Tavaborole *(Kerydin)* as a potent fungistatic agent against *Candida auris*"

The primary *in-vitro* spotting assay utilized for the initial assessment of the inhibitory effect of compounds identified by the *in-silico* screening strategy. Assays were carried out using RPMI-MOPS with 2% glucose and 2% agar plates with final drug concentration of 50µM, *Candida auris* isolates belonging clade I (AR389), clade II (CBS10913), clade III (AR383), clade IV (AR385), clade V (AR1097), *Candida albicans* (SC5314) and *Candida glabrata* (ATCC2001) were used. Spotting was performed by a robotic arm, following which plates were monitored over time for fungal growth. The following compounds were screened: Carboxin (mw: 235.30), Cladribine (mw: 285.69), Colchicene (mw: 399.43), Dasatinib (mw: 488.01), Enasidenib (mw: 473.38), Fostamatinib (mw: 580.46), Griseofulvin (mw: 352.77), Mupirocin (mw: 500.62), Pemetrexed (mw: 427.41), Pimecrolimus (mw: 810.45), Selinexor (mw: 443.31), Sulfinpyrazone (mw: 404.12), Tavaborole (mw: 151.93), Triclabendazole (mw: 359.65) and additionally AmB 2µg/mL, control RPMI-MOPS and control YPD plates.

1. Carboxin 24h 48h

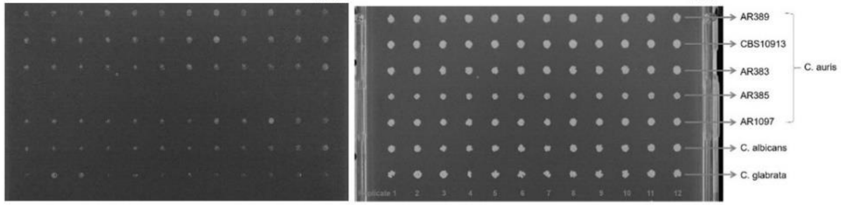


2. Cladribine 24h 48h

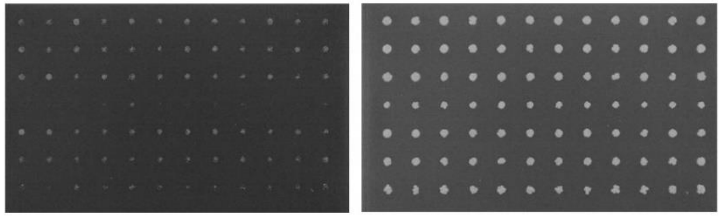

3. Colchicene 24h 48h

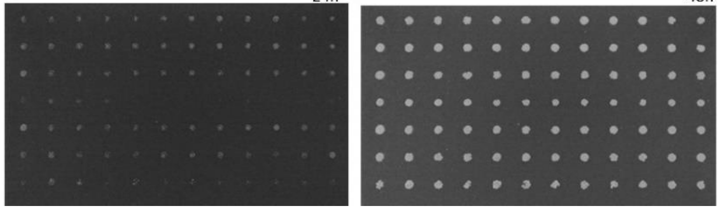

4. Dasatinib 24h 48h

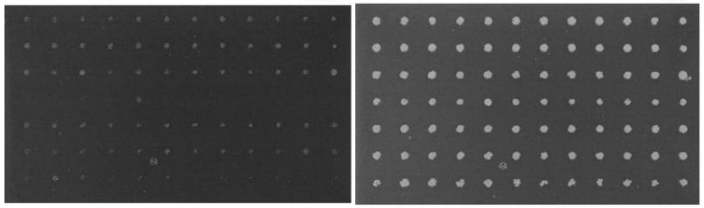

5. Enasidenib 24h 48h

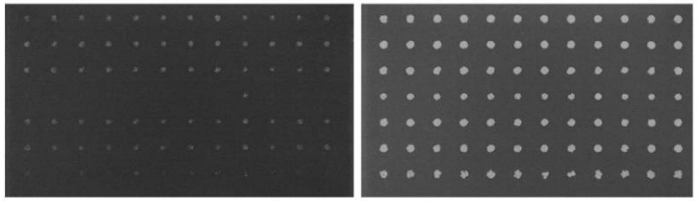

6. Fostamatinib 24h 48h

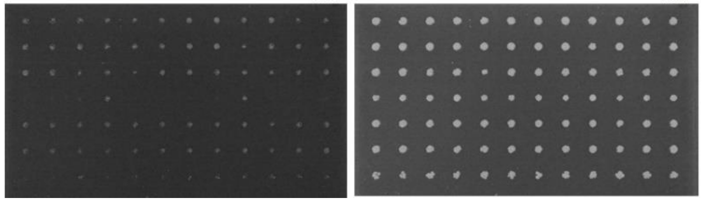

7. Griseofulvin

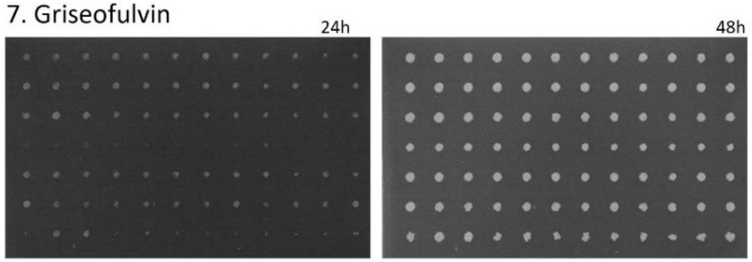


8. Mupirocin

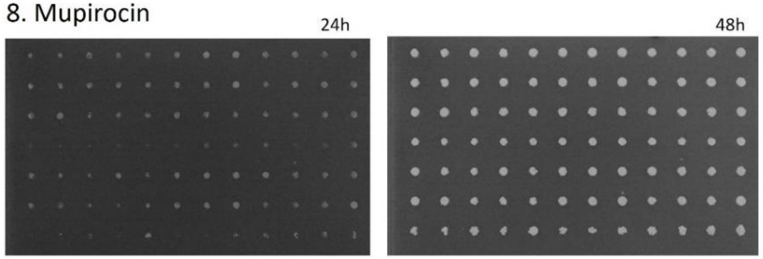


9. Pemetrexed

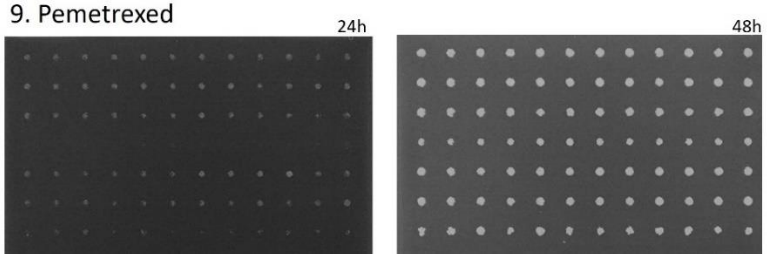


10. Pimecrolimus

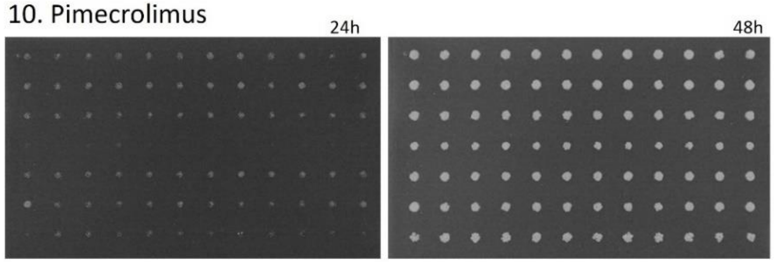


11. Selinexor

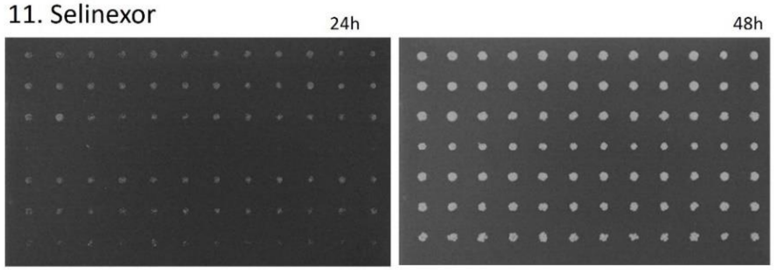


12. Sulfinpyrazone

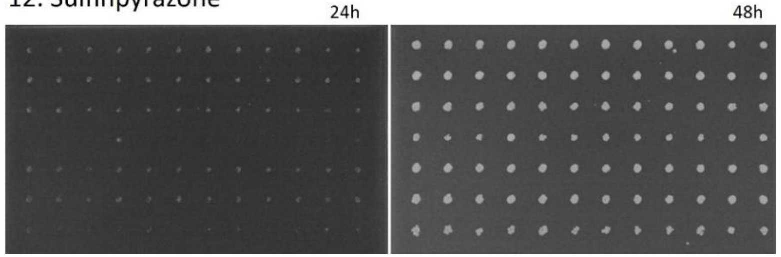

13. Tavaborole

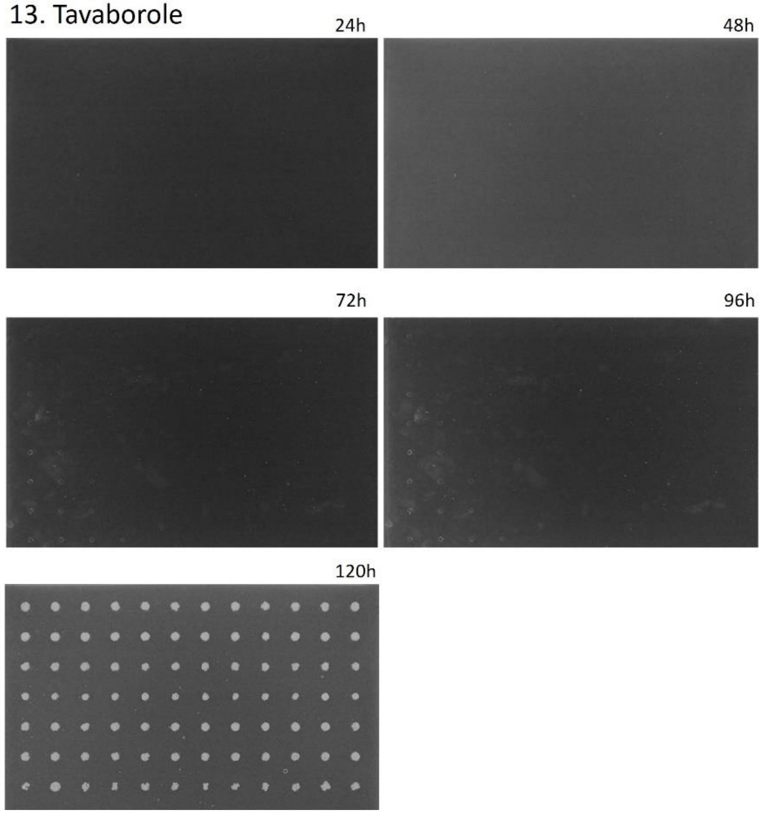


14. Triclabendazole

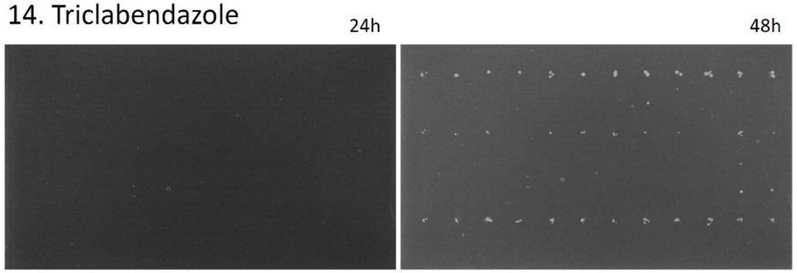


15. AmB

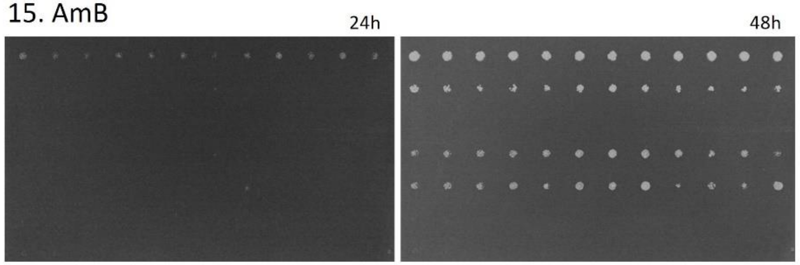


16. RPMI-MOPS control

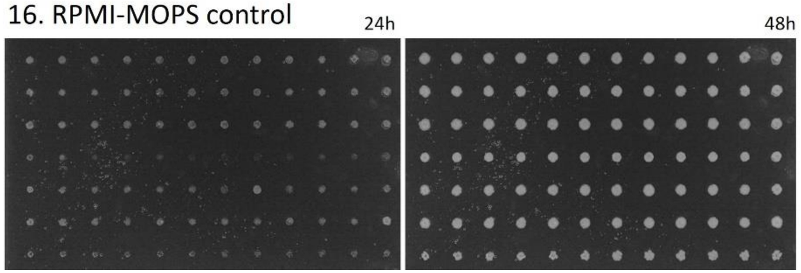


17. YPD control

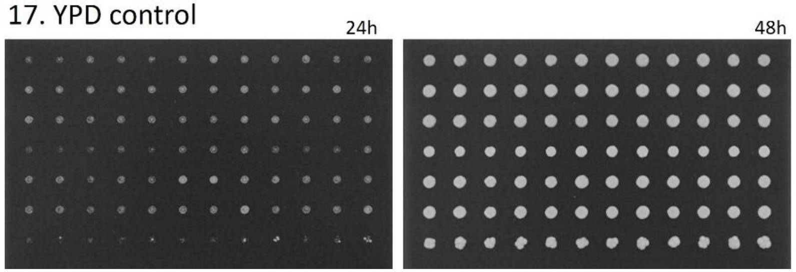
